# Prioritizing Combinational Drug Screening: A Ranking System for *In Vitro* Drug Combinations in Neurofibromatosis Type 1

**DOI:** 10.1101/2025.08.02.667047

**Authors:** Kangyi Zhou, Syeda Shehzad, Zaynab Ahmed, Olivia Zhao, Carolina Sapriza, Matthew Zamora, Ulisses Santamaria, Paul Zamora

## Abstract

Neurofibromatosis type 1 (NF1) is a genetic disorder characterized by benign tumors, including plexiform neurofibromas, which can be difficult to treat. Currently, only two FDA-approved therapies exist: selumetinib, approved for pediatric patients with inoperable tumors, and mirdametinib, approved for patients aged two and older with symptomatic peripheral neuropathy where surgical resection is not possible. These limited options highlight the urgent need for novel therapeutic strategies, including combination therapies and therapies applicable to adult populations.

In this study, we introduce the Composite Matrix Reduction Score (CMRS), a novel algorithm designed to evaluate the *in vitro* efficacy of drug combinations for NF1-related plexiform neurofibromas. Using a high-throughput 6×6 combinatorial matrix, we screened three cell lines: ipnNF95.11c (*NF1+/-*, non-tumor reference), and two *NF1-/-* tumor lines: ipNF05.5mc and ipNF95.6. Cell viability responses to drug combinations were normalized to vehicle controls, and combination effects were compared to single-agent responses. Tumor-to-non-tumor response ratios were aggregated to generate a composite ranking for each drug pair.

Our results show that certain drug combinations outperformed single agents in reducing tumor cell viability, consistent with findings in other cancers. A focused analysis on selumetinib combinations supported the CMRS algorithm and identified potential synergistic partners that may surpass a single-agent therapy, highlighting candidates for continued investigation.

CMRS provides a scalable, standardized framework for prioritizing drug combinations in NF1 and potentially other cancers. By integrating multi-cell line analysis, this approach enhances the identification of promising therapeutic candidates and mechanisms of action for further preclinical development.

## 1. Introduction

Neurofibromatosis type 1 (NF1) is a rare, chronic genetic disorder characterized by heterozygous germline NF1 gene mutations. Pathogenic variants of the NF1 gene can lead to various neurofibromas involving Schwann cells, including plexiform neurofibromas (1). Homozygous NF1 mutation is typically associated with loss of function of the neurofibromin protein, the NF1 gene product. Patients with plexiform neurofibromas have an increased risk of malignancy, 10% to 15% of NF1-affected individuals develop malignant peripheral nerve sheath tumors (MPNSTs) (2). There are two FDA-approved drugs used in the treatment of neurofibromatosis type 1 (NF1)-associated plexiform neurofibromas: selumetinib and mirdametinib (3-6). Both selumetinib and mirdametinib are inhibitors of MEK (mitogen-activated protein kinase kinase; MAPKK). Selumetinib is approved for certain pediatric patients, while mirdametinib (PD 0325901) was recently approved for the treatment of certain adult and pediatric patients. Nonetheless, some patients do not respond or only partially respond to treatment (4, 7), and protracted treatment may result in drug resistance in patients who have a positive initial response. This highlights the need to develop additional drugs and/or drug combinations that could be used in the treatment of NF1 syndromes.

Given that only two single-agent treatments are currently FDA-approved, combination therapies may represent a promising approach for overcoming some of the limitations of single-agent treatments (8, 9). Specifically, in the context of cancer therapy, combination treatments have shown significantly improved efficacy for specific types of cancer, such as acute lymphoblastic leukemia and Hodgkin’s lymphoma (9, 10). For many cancers, combination therapies can improve overall survival and progression-free survival (9). A similar approach is actively being studied in vitro and in vivo in NF1-related tumors (11, 12, 13).

While combination therapies offer considerable promise, a major methodological challenge persists in how best to evaluate and prioritize them. High-throughput screening (HTS) has emerged as a powerful tool in cancer drug development, enabling the rapid assessment of numerous drug candidates on cancer cells to identify those with the greatest therapeutic potential. However, there are limitations in accurately prioritizing these candidates based on the data generated. Traditionally, this prioritization has relied on metrics like AC50 (the concentration needed to achieve half-maximal activity) and the area under the curve (AUC) of dose-response curves (14-16). While AC50 can be useful, it has notable limitations—especially in high-throughput screens with limited dose points. Sparse sampling often yields partial dose-response curves that limit the accuracy of AC50 estimates. Additionally, even when there is adequate sampling, certain drugs can exhibit shallow curves, also skewing AC50 interpretation. Similarly, AUC provides a different measure of drug efficacy but can obscure critical differences in drug responses (8).

To address this methodological gap, we have developed the “Composite Matrix Reduction Score” (CMRS) as applied to a 6×6 drug-combination matrix format. Having just six concentration samples limits dose response coverage, especially considering that one of the six concentration levels is a zero-concentration control. This eliminates logistic curve analysis, as EC50 values are artificially skewed from sampling bias. Regardless, the CMRS can bring in useful insights in similar resource-constrained preliminary screens.

## 2. Methods

### 2.1 Data

High-throughput screening data used in the current evaluation were generated by Ferrer et. al. (18), in which cell lines generated from patients with NF1 syndrome were challenged with single and combined approved and experimental drugs in a cell proliferation assay. The NF1+/− non-tumor cell line ipnNF95.11c was derived from normal peripheral nerve tissue of a patient with NF1 syndrome. The NF1-/-cell lines (ipNF95.6 and ipNF05.5) were derived from plexiform neurofibromas. All data were accessed via the Synapse Portal (19, available at https://www.synapse.org/Synapse:syn5611796).

Each 6×6 matrix represents two compounds tested in a series of five 5-fold dilutions – the rows of the matrix being one compound doses, and the columns being the other compound doses. The sixth final row or column was a no-drug baseline, thus resulting in a total of 25 unique drug/dosage combinations for each test plate, one row of a single compound, and one column of the other respective single compound, see Figure 1A. The drugs/compounds were part of the Mechanism Interrogation PlatE (MIPE 4.0) library. The dataset included 780 drug combination matrices per cell line, each identified by a unique block identifier (BlockID). Each BlockID corresponds to a distinct pair of drugs tested across a range of concentrations. This layout forms a 6×6 drug combination matrix, which is the basis for the CMRS approach in Figure 1.

**Fig. 1.**
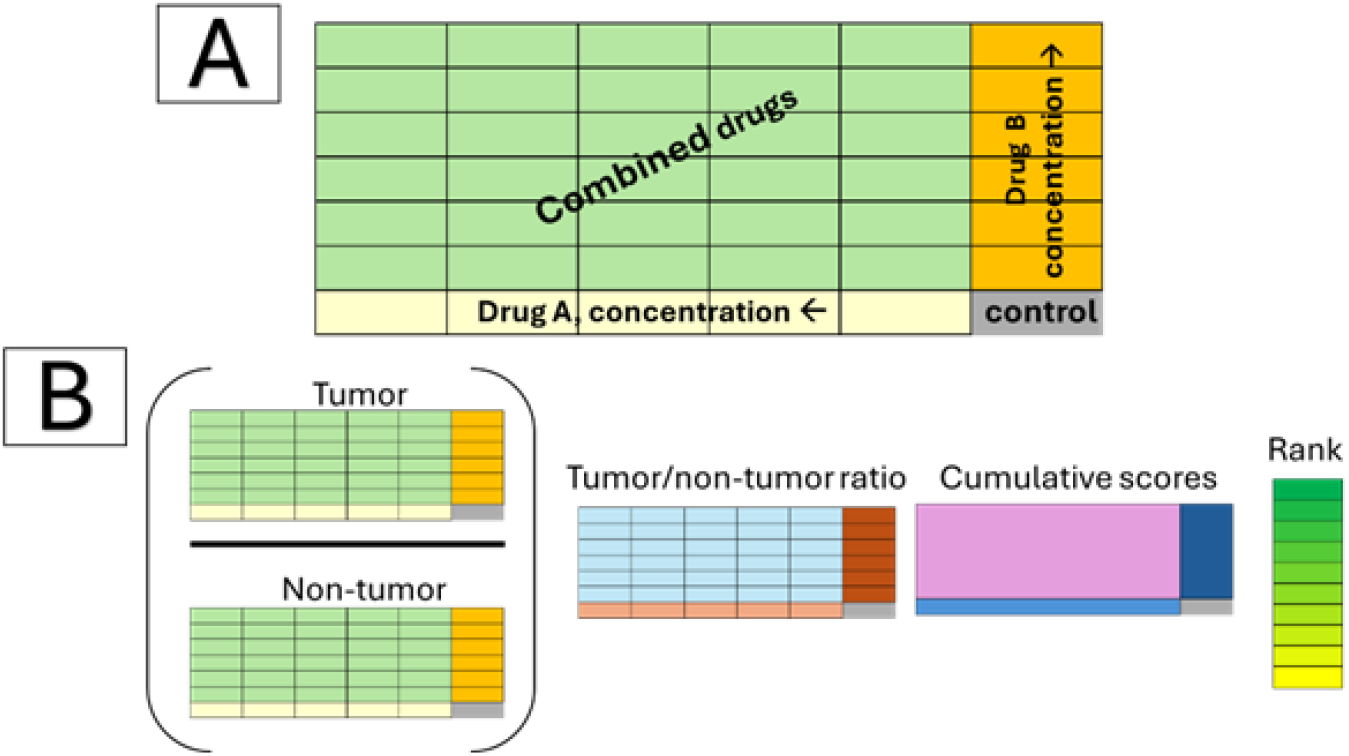
CMRS Flow Diagram. **A.)** Representation of the 6×6 combinational drug matrix. Drug A is represented in light yellow as a single agent zone (i.e. drug B concentration is zero). Likewise, a single agent zone for drug B in orange. Lastly a combination zone in green shows where drug A and drug B have additive concentrations (concentrations above zero). The solvent-only control (grey) serves as the baseline for normalization. **B.)** After a normalization calculation, implied to occur prior to the ratio shown here, the tumor and non-tumor response matrices (bracketed matrices) are used to determine a single ratio matrix, with the tumor/non-tumor response ratios (light blue) for each drug used (dark brown and light brown each respective to a single drug). These ratios are assigned scores between 0-6 based on predefined ranges to create a matrix of cumulative scores. These scores were then aggregating in each region – purple (combination therapy region), and blue and dark blue (respective single agent regions) – generating a final cumulative score per drug combination for each of the three regions. Final cumulative scores closer to zero indicate that the reference had a more dominant effect than the test compounds. Higher scores show selective effect in tumor versus non-tumor samples.

### 2.2 Composite Matrix Reduction Score

Figure 1 presents a flow diagram of the process used to generate the Composite Matrix Reduction Score (CMRS).

Each matrix was normalized to the response at the solvent-only control well (Figure 1A, gray control zone; see Equation 1)—where both drugs were at zero concentration. This calculation standardizes the data to reflect a percentage relative to the solvent-only control response. The n value is the response at a given position in the matrix, and the control value is the specific value where both compound concentrations are zero.

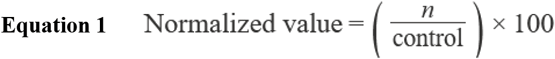

Each drug combination matrix was normalized independently for both the reference and test cell lines. After normalization, a unified ratio matrix was generated for each drug pair by dividing the normalized values of the test cell line by the corresponding values in the reference cell line at each matrix position. These intermediate ratios provide a comparative measure of drug response between cell types. Ratios less than 1 indicate increased sensitivity in the tumor (test) cell line relative to the non-tumor (reference) line, whereas ratios greater than 1 suggest relative resistance in the tumor line. A ratio near 1 reflects comparable responses in both cell lines.

To simplify the interpretation of these ratios, a scoring system was applied to each cell based on specified ranges (see Table 1). The method of choosing the score bins involved looking at the distribution of all ratios over a bell curve of all samples. Break-points were selected to create near-evenly distributed bins while preserving meaningful distinctions in confidence levels. These thresholds may require adjustment depending on the specific characteristics of each assay.

**Table 1.** Scoring ranges applied to the intermediate ratio matrix to quantify the relative impact of the test cell line versus the reference cell line. Scoring ranges divide the ratios into seven intervals, with scores from 0 to 6 assigned based on the magnitude of the ratio. Lower ratios (e.g., ≤0.4) correspond to higher scores, indicating greater tumor cell sensitivity relative to reference. Higher ratios (e.g., >0.98) correspond to lower scores, reflecting reduced sensitivity. Priority scores are given to ratios that show clear selective behavior between tumor and non-tumor samples.

| Ratio Range | Score |
| --- | --- |
| $\leq 0.4$ | 6 |
| $> 0.4$ and $\leq 0.5$ | 5 |
| $> 0.5$ and $\leq 0.6$ | 4 |
| $> 0.6$ and $\leq 0.8$ | 3 |
| $> 0.8$ and $\leq 0.9$ | 2 |
| $> 0.9$ and $\leq 0.98$ | 1 |
| $> 0.98$ | 0 |

In the CMRS approach, seven ratio ranges were used to assign scores as shown in Table 1. Allocating ratio ranges into specific groups makes it easier to give them their corresponding scores and to analyze them further. To identify the specific ratio ranges, a histogram of the number of drug combos for each specific ratio range was created to visualize the distribution (see Figure 2). The histogram shows that there are less than 500 combinations for ratios of 0.4 and below. Ratios between 0.4 and 0.5 pertain to around 500 combinations. Between 500-1000 combinations demonstrate a ratio between 0.5 and 0.6. There are around 1000 combinations for the ratio range of 0.6 to 0.8. Around 1500 combinations demonstrate a ratio between 0.8 and 0.9. The ratios between 0.9 and 0.98 pertain to slightly more than 1500 combinations. The number of combinations relating to the ratio range of 0.98 and above is shown to increase and then decrease, with a spike of almost 2500 combinations for the specific ranges of (0.99, 1.03] and ending at almost 0 combinations at the range of (2.14, 2.18]. Seven ratio categories were used due to the number of combinations fluctuating throughout the ratio ranges. The selection of the ratio ranges was established based on the shape of the histogram curve.

**Fig. 2.**
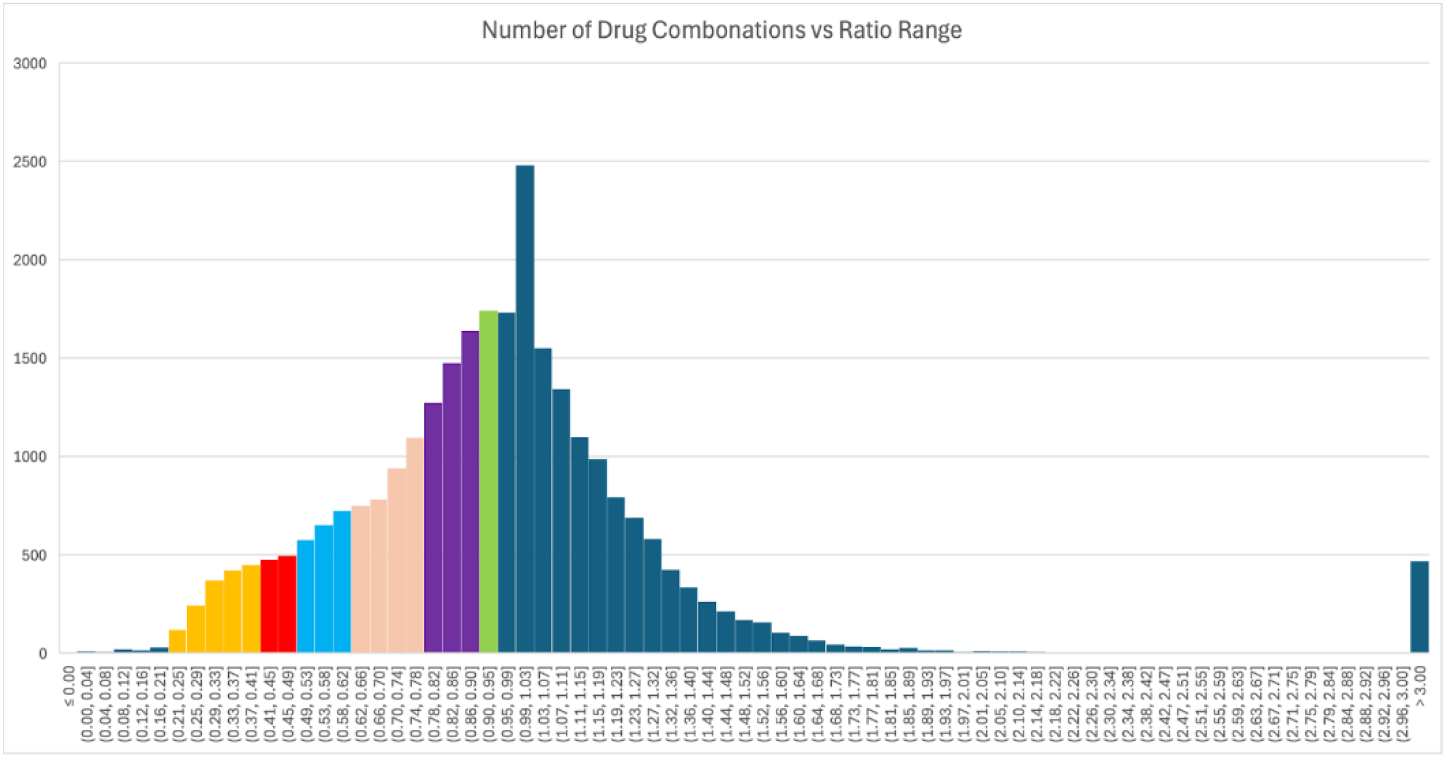
Histogram showing the distribution of drug combinations across ratio ranges. The x-axis is the ratio ranges of the drug combinations in bins, and the y-axis is the frequency of drug combinations within each range. Each colored group was a balance of evenly distributing scores and based on the shape segments of the histogram curve.

For each score matrix, a cumulative score was determined for all combination values within that block. Separately for each block, each set of single drug values was used to generate a cumulative score for each separate drug. (See Figure 3).

**Fig. 3.**
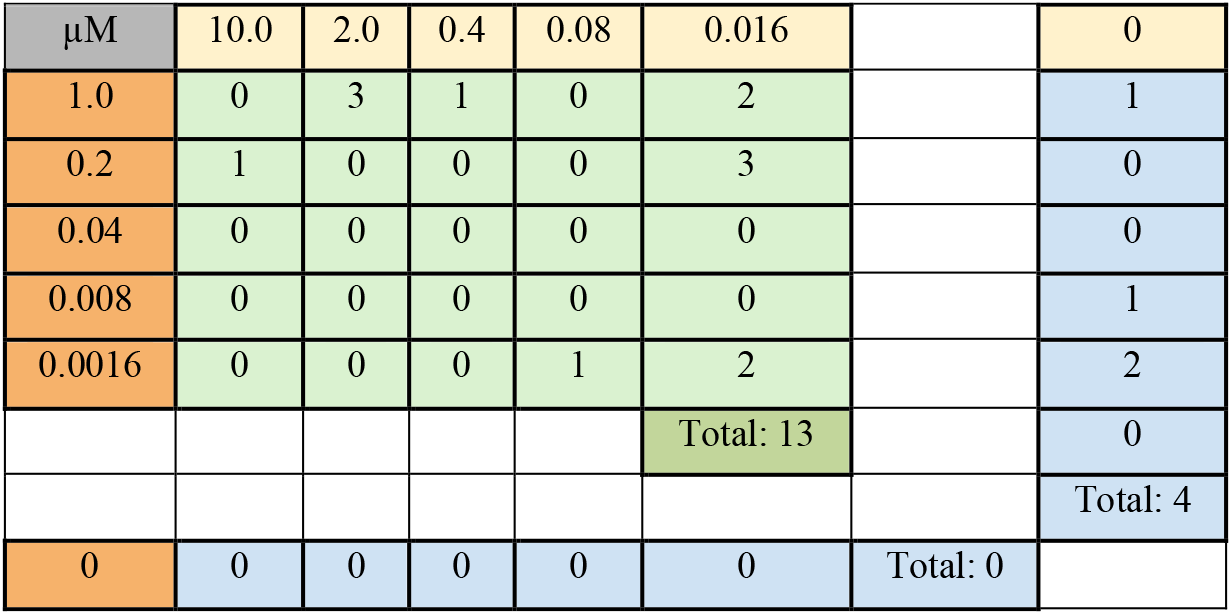
A conceptual mockup of scoring done by zones. Drug A in light yellow, and Drug B in orange have a distribution of concentrations. Three scoring zones are created: the combinational zone (green), vertical single-agent zone (orange), and horizontal single-agent zone (yel-low). Some forms of analysis compare single agent versus combinational therapy; in that case, the single-agent zones are considered equivalent and mergeable.

The three scoring zones identified are the combinational zone, the vertical singleagent zone, and the horizontal single-agent zone. These zones are visually represented in Figure 3 to clarify their definitions. In an analysis comparison of single agent effects versus combinational agent effects, both horizontal and vertical zones were merged and considered a unified zone.

Subsequently, the cumulative scores within the combinational zone across all drug combinations (BlockIDs) were calculated and ranked in descending order to assess their relative influence. Similarly, for the single-agent zones, the scores for each row and column were summed and ranked from highest to lowest across drug combinations to evaluate their individual contributions to the overall drug efficacy.

### 2.3 Combined Distribution Analysis

After generating ranked drug combination scores based on normalized tumor-to-reference cell line ratios, we conducted a comparative analysis to evaluate how closely the rankings of each combination matched between the tumor and reference cell lines. This analysis produced a histogram of absolute differences in composite scores for each drug combination, quantifying the magnitude of variation across cell lines (see Figure 7).

To focus on the most relevant drug combinations, the dataset was filtered using two criteria. First, only combinations with a magnitude difference of ≤10 were retained, ensuring a baseline level of similarity. Second, to prioritize combinations with relatively high efficacy, a composite ranking filter was applied. For each BlockID/drug combination, the rank position number of the reference and test cell lines were summed, and only combinations with a total rank ≤100 was included—corresponding to individual ranks of approximately 50 or better. This dual-filtering strategy balanced response similarity with therapeutic potential.

From the filtered dataset, two summary tables were generated: one highlighting the top five drug combinations with the greatest magnitude differences, and another showing the five combinations with the least variation. Each table includes the BlockID, row and column drug names, mechanisms of action for both compounds, individual ranks and composite scores per cell line, and the summed composite score for each combination.

## 3 Results

### 3.1 Single compound responses

To assess the overall benefit of a combinatorial drug approach when compared to single agent-only approaches, we plotted the distribution of responses of each cell line to the maximum dose of each drug separately and of each combination, presented as a percentage of cell viability of the solvent-only response (see Figure 4). Single-agent responses (orange bar) generally ranged between 50% and 60%. In contrast, the highest concentration of combined agents resulted in a median viability score of 15% to 35%, demonstrating a substantially greater effect than the single agents alone. The same trend is exemplified in Figure 6, where the combination showed lower cell survival of tumor cells.

**Fig. 4.**
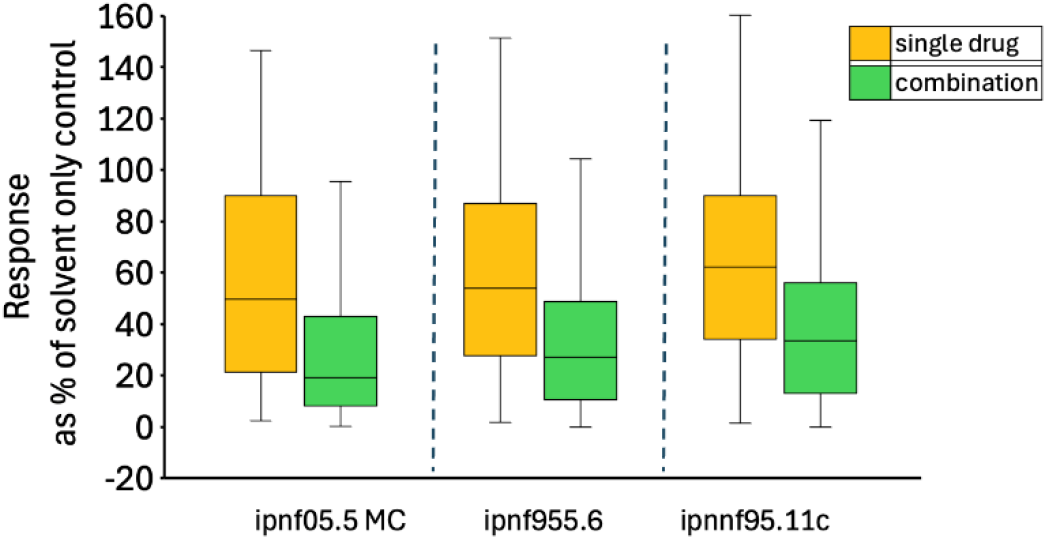
Demonstration of the maximum effect observed when looking exclusively at single drug zones, versus combinational therapy zones. The figure demonstrates the single drug zones in orange and is inclusive of both the horizontal and vertical single-agent zones. The green are the combinational drug zones. For each drug combination, only the lowest observed cell viability value from the single-agent and combination zones was used per cell line to represent the maximum effect. The distribution of the observed effect size across all drug combinations is then shown in the box and whisker plots, for each of three cell lines: ipNF05.5MC (*NF1-/-*, tumor), ipNF95.6 (*NF1-/-*, tumor), ipnNF95.11c (*NF1+/-*, non-tumor).

Despite the overall trend in Figure 4, in some cases, a single compound demonstrated greater efficacy than its combination counterpart. For example, BlockID 32 tested the MEK inhibitor PD 0325901 in combination with the VEGFR-2 inhibitor Cabozantinib. Plexiform neurofibromas are known to be sensitive to MEK inhibition (20), but not to VEGFR-2 inhibition, so Cabozantinib alone was expected to have minimal impact on cell viability. Consistent with this, Cabozantinib at 10 μM (the highest tested concentration) reduced viability in the non-tumor ipNF95.11c cell line to 38%, and in tumor cell line ipNF05.5mc to 41%, yielding a tumor-to-control response ratio of 1.08—indicating no increased tumor sensitivity.

In contrast, PD 0325901 at concentrations up to 5 μM had no effect on the non-tumor cells (100% viability), but reduced tumor cell viability to an average of 86%, corresponding to a ratio of 0.86 and confirming tumor-specific sensitivity. When combined, PD 0325901 and Cabozantinib reduced viability to 29% in tumor cells and 31% in non-tumor cells, resulting in a ratio of 1.06. This suggests that the combination did not enhance tumor-specific efficacy beyond what was observed with PD 0325901 alone.

The pattern for PD 0325901/Cabozantinib suggests that the lowest response in the combination was driven by the compound with the highest toxicity when used alone; in other words, the reduction of viability overall (in both control and tumor cell lines) suggest some combinations rank low due to additive toxicity. This observation is consistent with independent drug action (IDA), which hypothesizes that the expected effect of a combination of non-interacting drugs is simply the effect of the single most effective drug in the combination and this concept has been developed into a combination drug predictive tool (IDA Combo) (21). A key goal of this study is to identify any additional response patterns that may be more clinically useful.

### 3.2 Ranking Results

The effectiveness or non-effectiveness of the first twenty and last twenty drug combinations respectively were systematically interpreted relative to a literature review of the drugs when used as both single agents as well as combinational agents, if applicable. Drugs were validated for their respective placement on the generated ranked list. Consideration was given for known signaling pathways in NF1. Each ranked item was assigned a number for literature support at the time of the analysis in relation to its rank: 1-10 with a score of 1 being the most supported and 10 being the least supported. These assigned scores served as a qualitative check to validate that the algorithm-generated ranking aligned with existing literature on NF1-relevant drug mechanisms.

Table 2 illustrates the respective CMRS combination scores from the ipNF05.5mc test cell line and the ipnNF95.11c reference line. The highest ranked drug combination tested in the ipNF05.5mc cell line was the Heat Shock Protein 90 (HSP90) inhibitor Alvespimycin with the topoisomerase inhibitor Topotecan. Despite Alvespimycin and Topotecan ranking highest by CMRS score, combinations with Topotecan were excluded from further discussion due to its limited clinical application related to mutagenic effects that could potentially lead to malignancies (22). Alvespimycin was a top contributor to the list, as one of the drugs in six of the top ten scoring drug combinations. Previous research validates Alvespimycin’s frequent appearance as it is a strong HSP90 inhibitor in combinational treatments (23). The second highest combination in ipNF05.5mc was Panobinostat and Ganetespib which are a histone deacetylase (HDAC) inhibitor and a HSP90 inhibitor, respectively. An HDAC inhibitor was found in four of the top ten drug combinations. In these drug combinations the cell response was consistent with the independent drug action model. Carfilzomib, a proteasome inhibitor, appeared as one component of a combination in Table 2 a total of nine times suggesting a potential, if currently unclear, role as a drug combination partner. Selumetinib’s role in section 4.4 is further discussed.

**Table 2.**
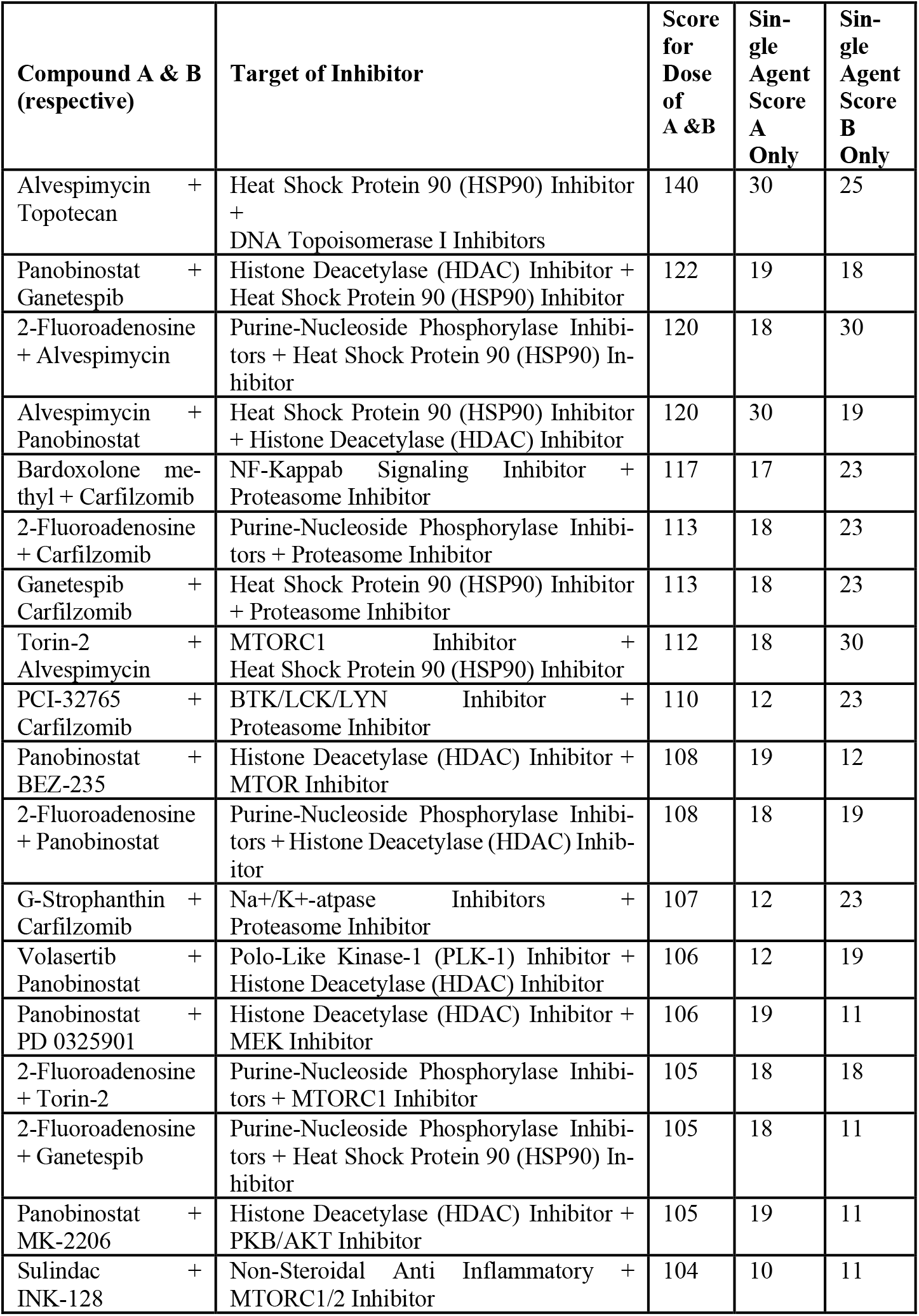
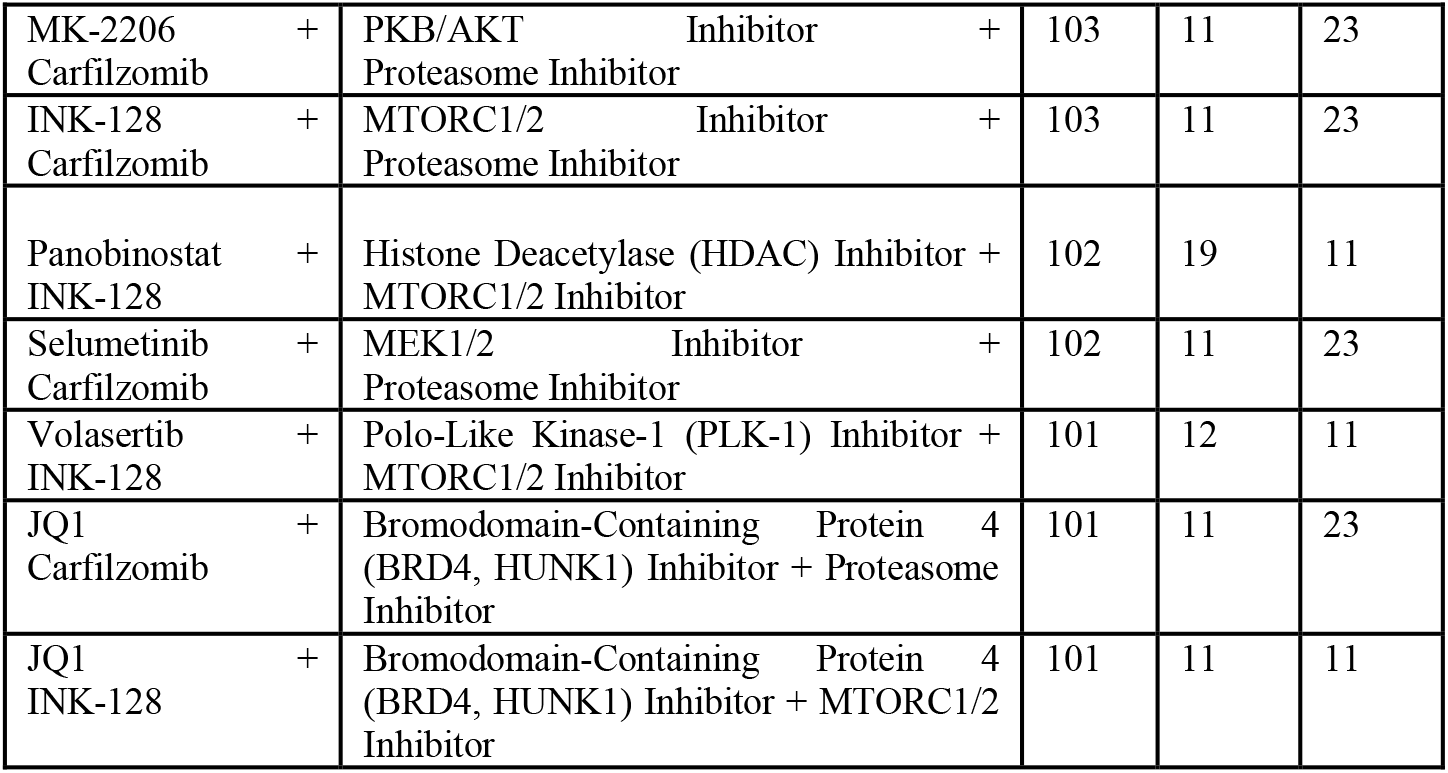
Summary of the top-ranked drug combinations tested relative to the ipNF05.5mc test cell line and the ipnNF95.11c reference line at a variety of concentrations, including their mechanisms of action, combinational scores, and individual scores for each single-agent drug.

The graphs shown in Figure 5 demonstrate two cell lines, ipnNF95.11c (non-tumor) and ipNF05.5mc (tumor). The blue bars represent a concentration gradient of drugs Panobinostat and Ganetespib, ranging from 0-5 (nM). The red bars show the percentage of cell survival when exposed to this drug combination, ranging from 0-120. The ipNF05.5mc cell lines demonstrate significantly less cell survival, mostly being under 20%, while the ipnNF95.11c cell line averages at around 50%. Both cell lines showed high cell survival at low concentrations with increasing concentrations generally demonstrating less cell survival although with variations in the ipnNF95.11c cell line. The comparison of the responses of the two cell lines, ipnNF95.11c (a non-tumor nerve tissue cell line) and ipNF05.5mc (a benign plexiform neurofibroma cell line) in Figure 5 provides a representative example of drug efficacy across each cell line. This figure is particularly useful for analyzing reference-to-test cell line comparisons, while Figure 5 focuses on drug-to-drug interaction.

**Fig. 5.**
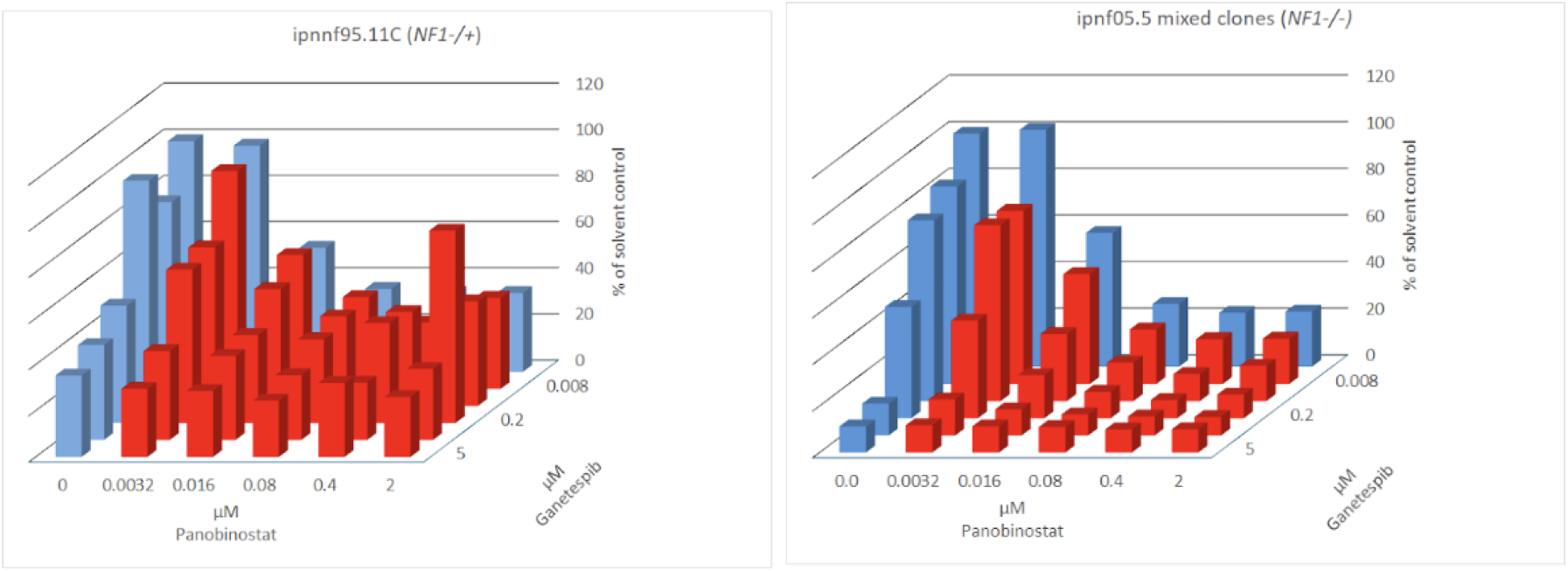
Comparative analysis of the non-tumor cell line ipnNF95.11c and the benign plexiform neurofibroma cell line ipNF05.5mc when exposed to Panobinostat and Ganetespib. Blue bars represent the concentration levels of the single-agent drugs used in combination, while red bars represent the percentage of cell survival. For the benign plexiform neurofibroma cell lines, significantly less cell survival can be observed compared to non-tumor nerve tissue cell lines.

**Fig. 6.**
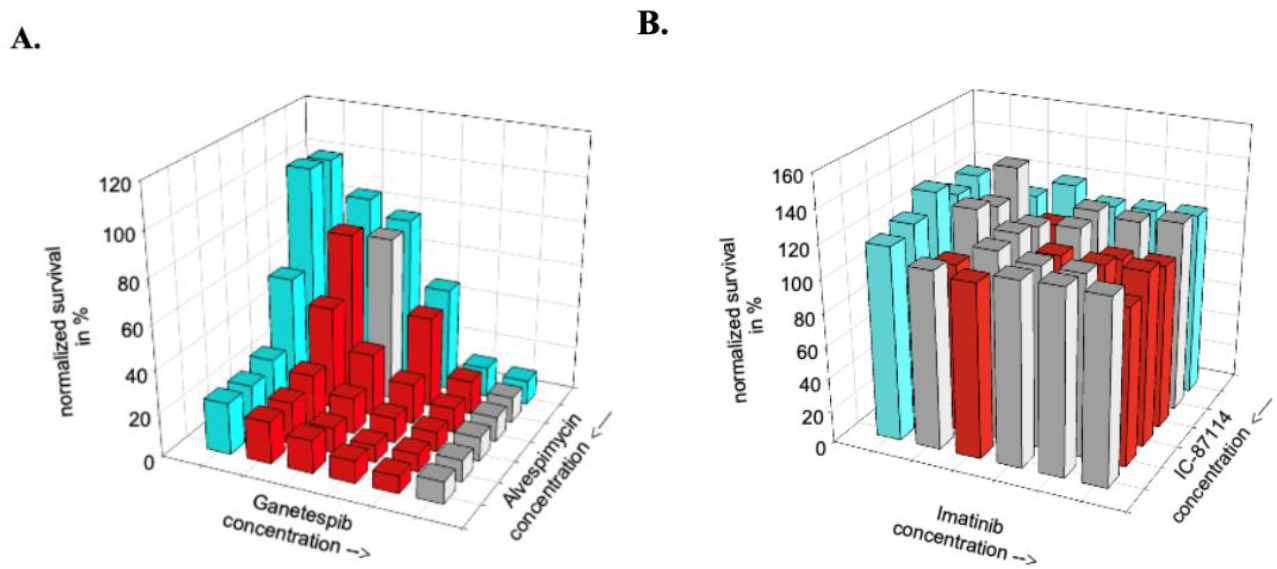
An alternative view of cell survival as a percent of solvent-only control, for two drug combinations in A and B. The blue bars show single-agent only zones, and the grey bars represent combinational zone values where the response was similar to (a less than 3%) the respective single agent zone. The red bars are combination compounds zones with responses greater than or equal to 3% different from respective single compounds. **A.)** 3D representation of the efficacy of both single-agent and combinational therapies involving Ganetespib and Alvespimycin, generally demonstrates that combinational therapies result in a greater reduction in cell survival compared to single-agent treatments. **B**.) Efficacy of the least effective drug combination as visualized in a 3D bar graph. There are fewer red bars, and all bars in the combinational zone remain predominantly above the control level (100% on the z-axis).

**Fig. 7.**
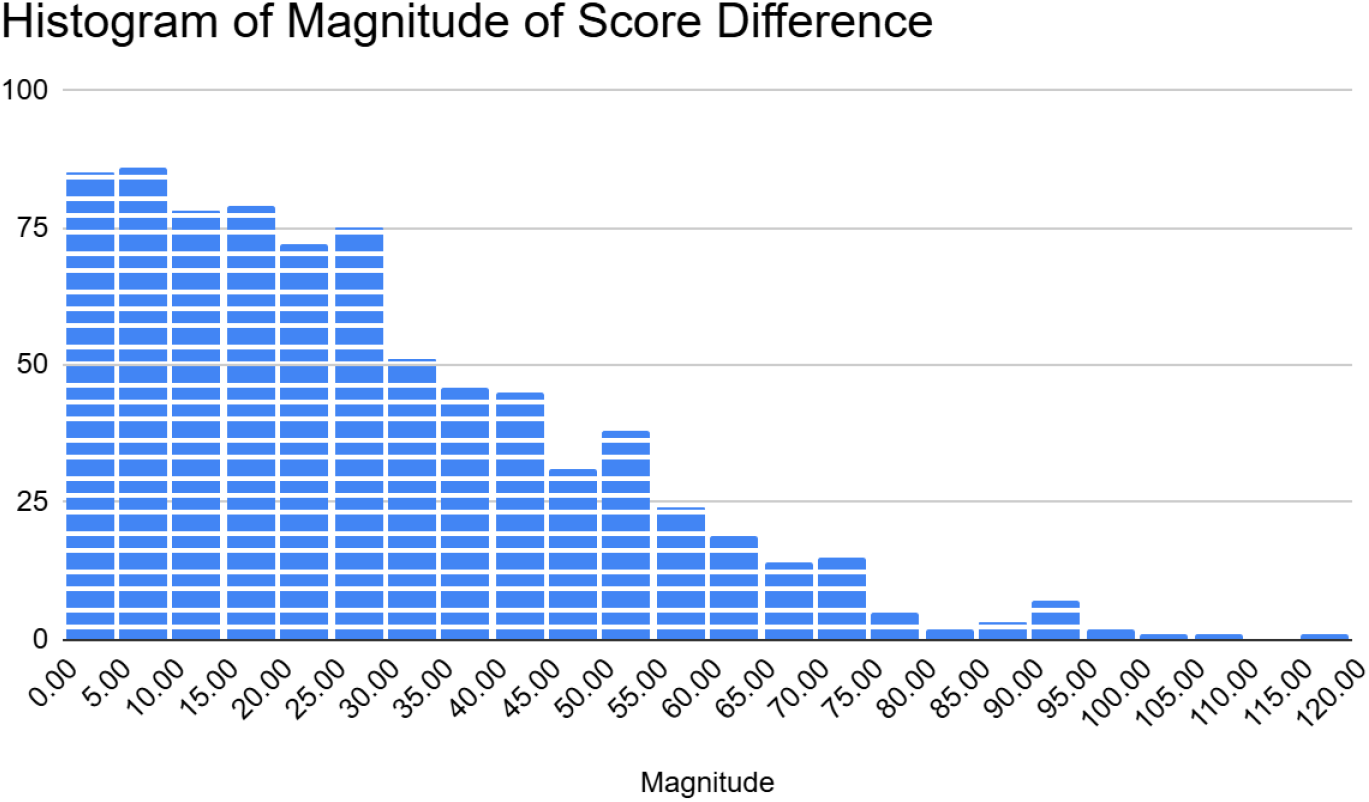
Histogram of the magnitude of score differences between cell test cell lines ipNF05.5mc and ipNF95.6. The x-axis is the magnitude of differences in total CMRS score for drug combinations and the y-axis is the frequency of drug combinations within each magnitude range. Lower magnitude values indicate greater similarity in drug response between the two cell lines. This is the same magnitude score difference as used in Table 4. The effects of many drug compounds were reproducible between test cell lines, but there are notable numbers that were cell-line specific.

While Figures 5 and 6 make spot checking between cell lines and drug combinations respectively an obvious activity, there are limitations when working with over 700 possible drug combinations. This makes it impractical to generate separate visualizations for each combination. The CMRS algorithm supports the early analysis by replacing the need for separate visualizations of each combinational therapy and eliminates the need to examine two separate graphs for every drug combination.

As shown in Figure 6, by creating a 3D bar graph illustrating the efficacy of different concentrates within the combinational therapy as well as single-agent therapies, the analysis provides a clear visualization of how varying drug levels influence cell survival, enabling a more precise evaluation of optimal therapeutic dosing. Furthermore, the trends observed reinforce the advantage of combinational therapies over single-agent treatments, as the latter consistently show higher survival rates across all tested scenarios.

An additional analysis was done with the ipNF95.6 (tumor) test cell line, and the ipnNF95.11c reference cell line (Table 3). The most frequently appearing drug among the various combinations was CHS-828, a nicotinamide phosphoribosyltransferase inhibitor (NAMPT). As of mid-2025, no nicotinamide phosphoribosyltransferase inhibitors have been FDA-approved for any indication and thus were not further examined. On the other hand, Alvespimycin plus Topotecan (previously discussed) was also in the top list and as part of a combination and so was carfilzomib.

**Table 3.**
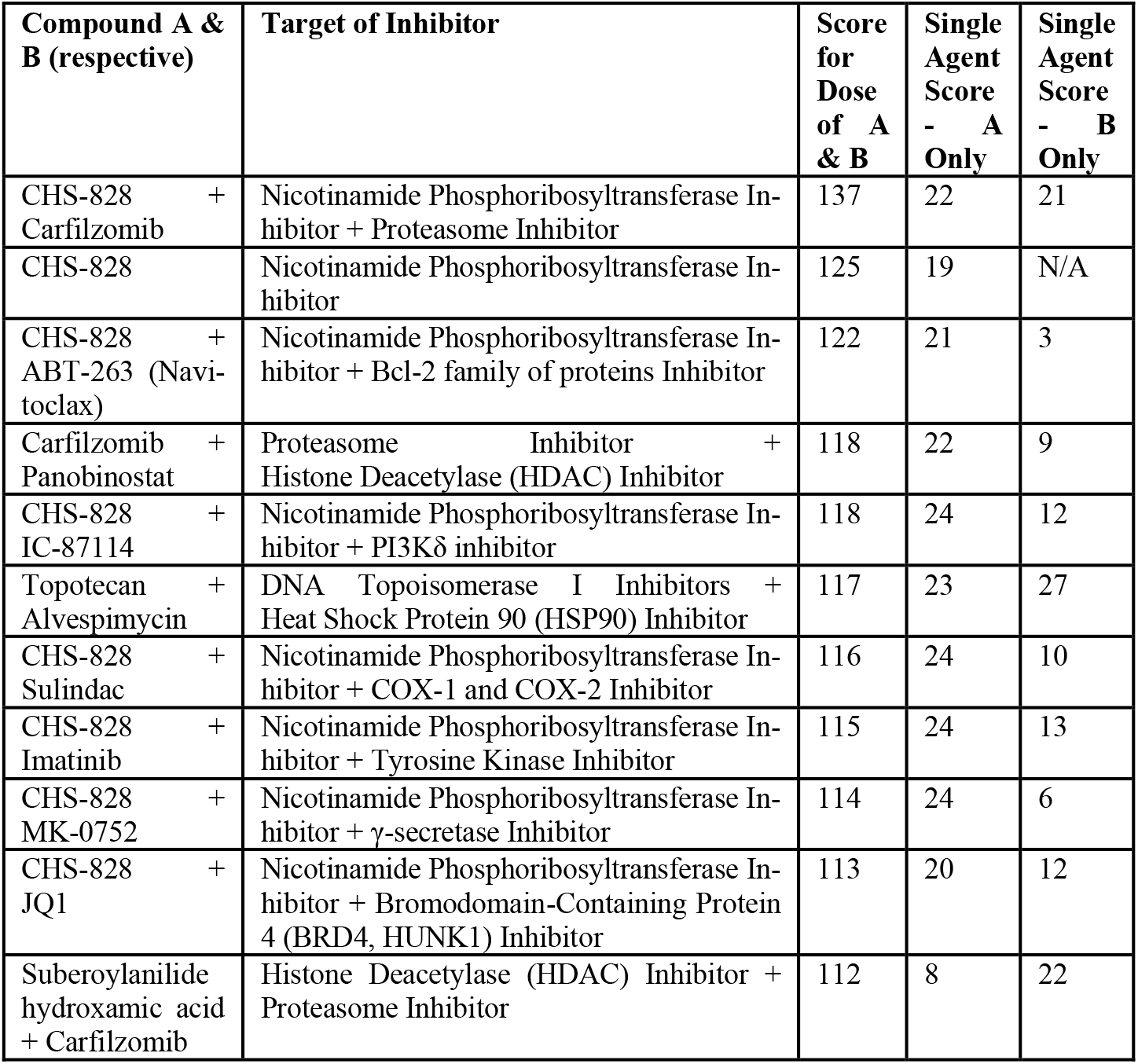
Summary of the top ten ranked drug combinations tested, including their mechanisms of action, combinational scores, and individual scores for each single-agent drug of the ipNF95.6 test cell line and the ipnNF95.11c reference cell line.

**Table 4.**
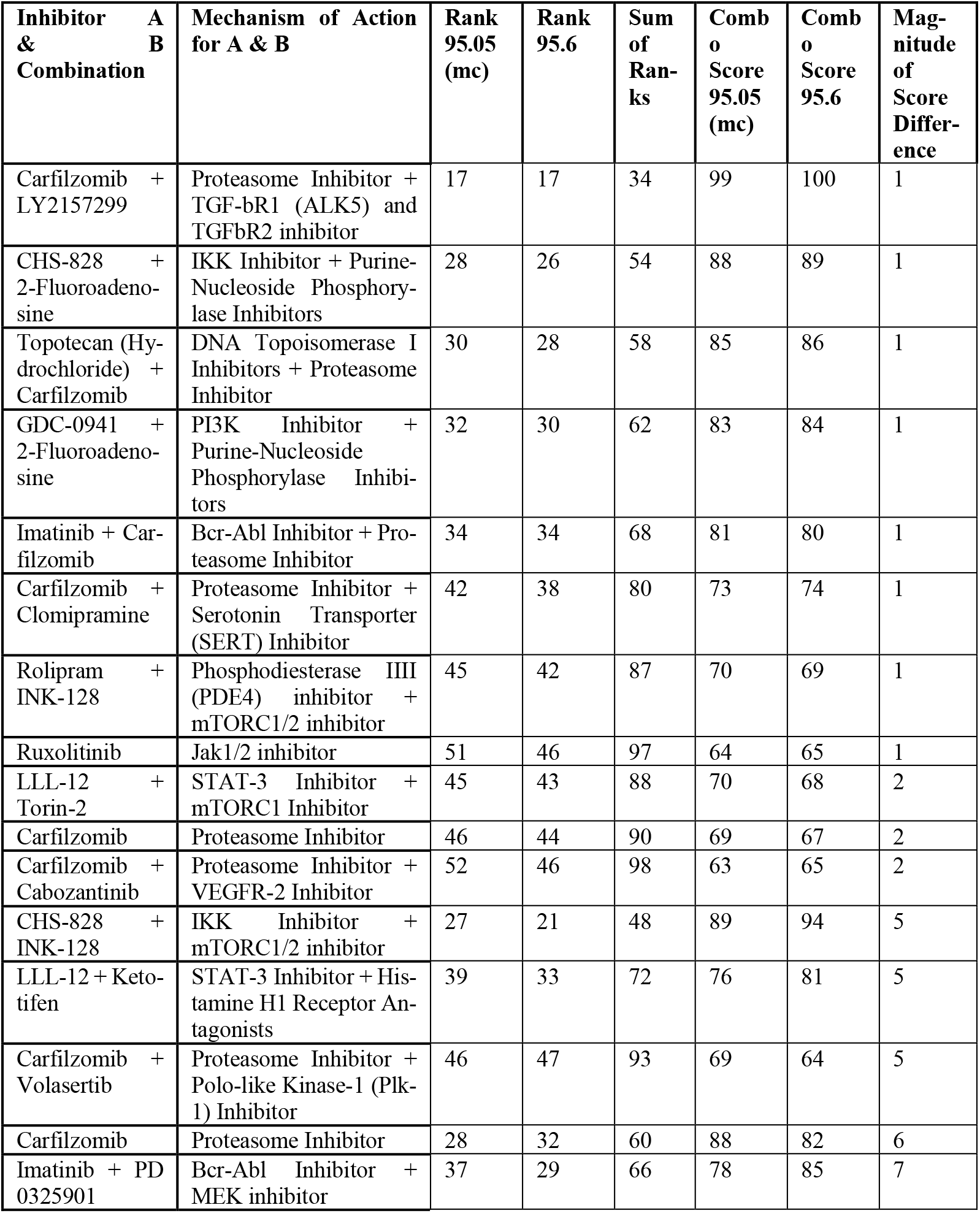
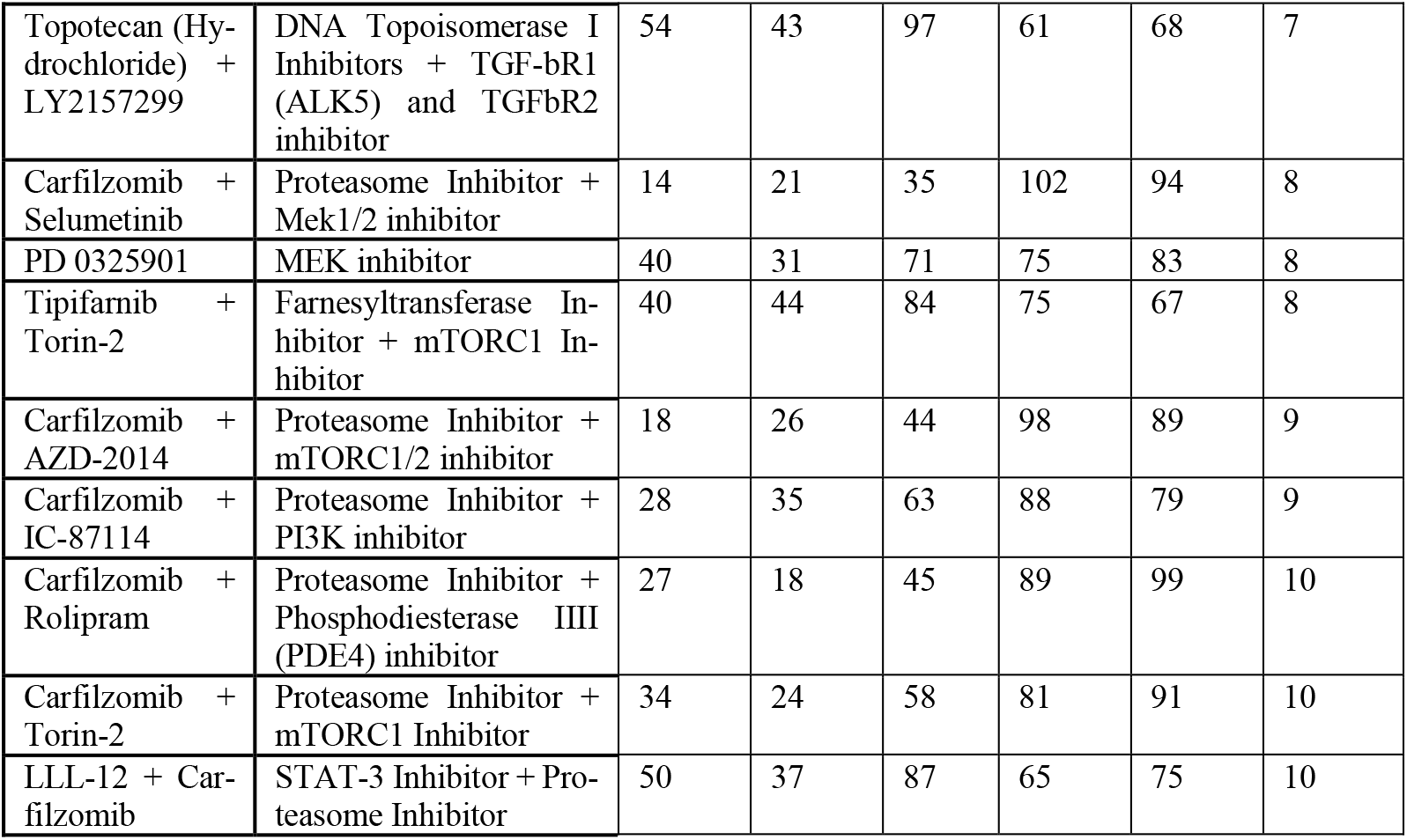
A cross-test cell line comparison, showing which drug combinations had repeatably observed effects. The table includes their respective rank position across all tested drug combinations, total CMRS scores, and the magnitude of score differences between the test lines. Lower magnitude values indicate a more similar drug response between the two cell lines, while higher values suggest greater variation in response.

| **Inhibitor A & Combination** | **Mechanism of Action for A & B** | **Rank 95.05 (mc)** | **Rank 95.6** | **Sum of Ranks** | **Comb o Score 95.05 (mc)** | **Comb o Score 95.6** | **Magnitude of Score Difference** |
| --- | --- | --- | --- | --- | --- | --- | --- |
| Carfilzomib + LY2157299 | Proteasome Inhibitor + TGF-βR1 (ALK5) and TGFβR2 inhibitor | 17 | 17 | 34 | 99 | 100 | 1 |
| CHS-828 + 2-Fluoroadenosine | IKK Inhibitor + Purine-Nucleoside Phosphorylase Inhibitors | 28 | 26 | 54 | 88 | 89 | 1 |
| Topotecan (Hydrochloride) + Carfilzomib | DNA Topoisomerase I Inhibitors + Proteasome Inhibitor | 30 | 28 | 58 | 85 | 86 | 1 |
| GDC-0941 + 2-Fluoroadenosine | PI3K Inhibitor + Purine-Nucleoside Phosphorylase Inhibitors | 32 | 30 | 62 | 83 | 84 | 1 |
| Imatinib + Carfilzomib | Bcr-Abl Inhibitor + Proteasome Inhibitor | 34 | 34 | 68 | 81 | 80 | 1 |
| Carfilzomib + Clomipramine | Proteasome Inhibitor + Serotonin Transporter (SERT) Inhibitor | 42 | 38 | 80 | 73 | 74 | 1 |
| Rolipram + INK-128 | Phosphodiesterase III (PDE4) inhibitor + mTORC1/2 inhibitor | 45 | 42 | 87 | 70 | 69 | 1 |
| Ruxolitinib | Jak1/2 inhibitor | 51 | 46 | 97 | 64 | 65 | 1 |
| LLL-12 + Torin-2 | STAT-3 Inhibitor + mTORC1 Inhibitor | 45 | 43 | 88 | 70 | 68 | 2 |
| Carfilzomib | Proteasome Inhibitor | 46 | 44 | 90 | 69 | 67 | 2 |
| Carfilzomib + Cabozantinib | Proteasome Inhibitor + VEGFR-2 Inhibitor | 52 | 46 | 98 | 63 | 65 | 2 |
| CHS-828 + INK-128 | IKK Inhibitor + mTORC1/2 inhibitor | 27 | 21 | 48 | 89 | 94 | 5 |
| LLL-12 + Ketotifen | STAT-3 Inhibitor + Histamine H1 Receptor Antagonists | 39 | 33 | 72 | 76 | 81 | 5 |
| Carfilzomib + Volasertib | Proteasome Inhibitor + Polo-like Kinase-1 (Plk-1) Inhibitor | 46 | 47 | 93 | 69 | 64 | 5 |
| Carfilzomib | Proteasome Inhibitor | 28 | 32 | 60 | 88 | 82 | 6 |
| Imatinib + PD 0325901 | Bcr-Abl Inhibitor + MEK inhibitor | 37 | 29 | 66 | 78 | 85 | 7 |
| Topotecan (Hydrochloride) + LY2157299 | DNA Topoisomerase I Inhibitors + TGF-βR1 (ALK5) and TGFβR2 inhibitor | 54 | 43 | 97 | 61 | 68 | 7 |
| Carfilzomib + Selumetinib | Proteasome Inhibitor + Mek1/2 inhibitor | 14 | 21 | 35 | 102 | 94 | 8 |
| PD 0325901 | MEK inhibitor | 40 | 31 | 71 | 75 | 83 | 8 |
| Tipifarnib + Torin-2 | Farnesyltransferase Inhibitor + mTORC1 Inhibitor | 40 | 44 | 84 | 75 | 67 | 8 |
| Carfilzomib + AZD-2014 | Proteasome Inhibitor + mTORC1/2 inhibitor | 18 | 26 | 44 | 98 | 89 | 9 |
| Carfilzomib + IC-87114 | Proteasome Inhibitor + PI3K inhibitor | 28 | 35 | 63 | 88 | 79 | 9 |
| Carfilzomib + Rolipram | Proteasome Inhibitor + Phosphodiesterase III (PDE4) inhibitor | 27 | 18 | 45 | 89 | 99 | 10 |
| Carfilzomib + Torin-2 | Proteasome Inhibitor + mTORC1 Inhibitor | 34 | 24 | 58 | 81 | 91 | 10 |
| LLL-12 + Carfilzomib | STAT-3 Inhibitor + Proteasome Inhibitor | 50 | 37 | 87 | 65 | 75 | 10 |

Beyond that, the highest-ranking drug pairing with selumetinib in the ipNF95.6 cell line was carfilzomib. Other pairs of interest pairing were CHS-828, LY2157299, Neratinib and Tipifarnib. Of interest for specific pairing with PD 0325901 in ipNF95.6 were CHS-828, Imatinib, LLL-12, GDC-0941 and AZD-2014.

The cellular responses to MEK inhibitor-based combinations generally aligned with the independent drug action (IDA) model from Huang et al. (21). For instance, selumetinib exhibited a dominant effect relative to carfilzomib, with enhanced activity observed in NF1-/-tumor cell lines compared to the NF1+/-reference line—consistent with expected genotype-specific sensitivity. A similar pattern was seen with PD 0325901 (mirdametinib), further supporting the role of MEK inhibition in these models. Overall, most drug combinations for both tumor cell lines produced outcomes broadly consistent with the IDA model, though some deviations were noted.

To assess the similarity and variability in drug responses between the tumor cell lines ipNF05.5mc and ipNF95.6, a histogram of magnitude differences was generated (see Table 4 and Figure 7).

Table 4 can demonstrate how combinations such as carfilzomib + LY2157299 and CHS-828 + 2-Fluoroadenosine exhibited minimal differences in composite scores (magnitude = 1), indicating highly similar responses across both cell lines. In contrast, combinations like carfilzomib + Torin-2 and LLL-12 + carfilzomib showed greater divergence (magnitude = 10), suggesting variability in efficacy between the two tumor models. The overall distribution of magnitude differences (Figure 7) reveals that most drug combinations clustered within lower ranges (≤20), with a gradual decline in frequency beginning around magnitudes of 30–35 and above.

## 4 Discussion

### 4.1 S’ and EC50 algorithm

In reviewing the 6×6 drug combinational data set by Ferrer et al. (18) used in this study, it was observed that dose-response curves for the single drug subsets of the data were incomplete and not full scale (see Figure 8). Limitations on dose coverage are common in combinational screening studies due to studies having non-linear costs for each additional concentration tested. In situations like these, the EC50 values may be ambiguous or inaccurate, and can be worsened when the data is restricted. We see in Figure 8 that the curves are abbreviated near the right descending shoulder. A similar ambiguousness can occur with AUC values or other methods that involve parameters based on the four parameter logistic model asymptotes and EC50. It has been suggested, in fact, that the relative EC50/IC50 should only be used if there are at least two assay concentrations beyond the lower and upper bend points (24). Unfortunately, this is not always a practical option in combinational high throughput screening.

**Fig. 8.**
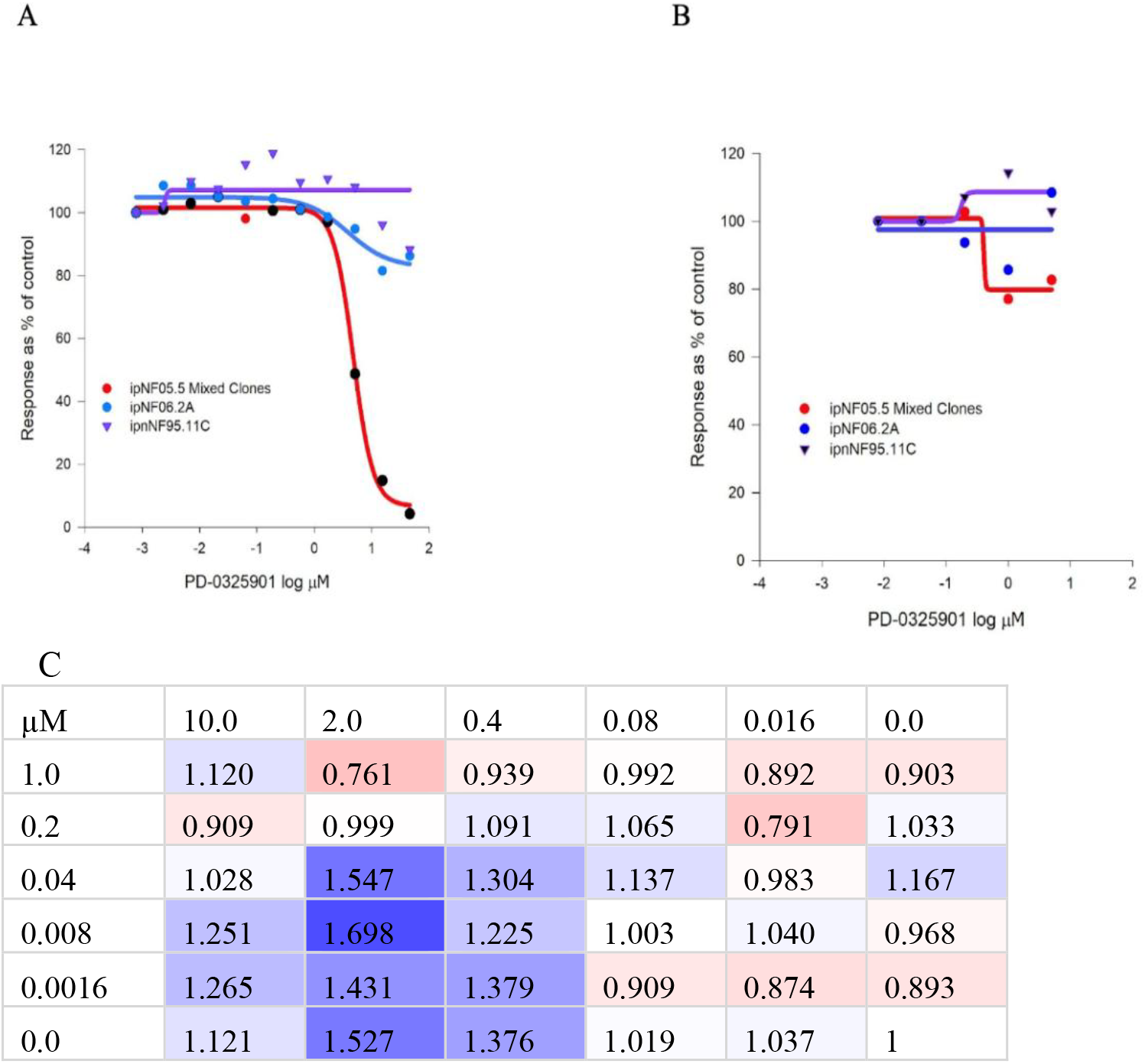
**A.)** Reference dose response curves from a related study (Zamora et al. (8)) using the same cell lines and compound library. Tumor cell lines (ipNF05.5 and ipNF05.2A) exhibited stronger dose-dependent responses, while the non-tumor reference line (ipNF95.11C) remained relatively stable across concentrations. **B.)** The corresponding dose response curve generated in the single-agent regions of the 6×6 combinational assay. The curves are not ideal and do not match. This would skew any analysis depending on EC50 or similar parameters. This is due to the model lacking data samples. **C.)** In contrast, the CMRS approach remains effective even with sparse data, highlighting combinations that outperform their respective single-agent treatments (i.e., where one compound is at zero concentration). In the normalized matrix, darker blue regions indicate favorable tumor-to-reference response ratios, while darker red regions reflect the opposite. Notably, a prominent blue block appears in the combination zone of AZD-2014 (row) and Cabozantinib (column), suggesting enhanced efficacy of the combination relative to either agent alone, warranting further investigation.

To help overcome these challenges, we developed a new algorithm without a strict reliance on EC50 and applied it to combination drug responses in plexiform neurofibroma cell lines. The benefits of the CMRS approach include its ability to analyze a large amount of data very quickly, present the combinational scores of each drug in the combination, and compare different combination therapies in a standardized manner, which can be visualized (see Figure 8C).

### 4.2 Notable Compounds

Certain patterns of the mechanisms of actions within drug combinations were observed. As seen in Table 4, carfilzomib consistently ranks highly in combination with other targeted therapies across both cell lines ipNF95.05 (mc) and ipNF95.6. Carfilzomib is a proteasome inhibitor that is usually used to treat patients with multiple myeloma, a type of cancer that primarily affects plasma cells. This drug selectively targets the proteasome, leading to the accumulation of misfolded proteins in cancer cells (30). The high levels of misfolded proteins will result in endoplasmic reticulum (ER) stress and the activation of the unfolded protein response, ultimately leading to apoptotic pathways (25). Since cancer cells have an exaggerated dependence on the proteasome, this allows carfilzomib to target tumor cells while minimizing the harmful side effects on the healthy cells (26).

Additionally, carfilzomib plays a role in inhibiting the nuclear factor-kappa B (NF-kB) signaling pathway by preventing the degradation of the IkB inhibitor, which suppresses the transcription of survival and proliferation genes in cancer cells (27). Other drugs, such as emetine and fluorosalan target the IkB inhibitor when used to inhibit the NF-kB pathway (28). This may lead to carfilzomib becoming a promising component of combinational therapy for NF1 patients, since NF1 mutations lead to hyperactive Ras signaling and dysregulated NF-kB activity (29). Additionally, NF1 tumors exhibit high proteasomal activity, making them vulnerable to the high ER stress and apoptotic effects that multiple myeloma cells experience when exposed to carfilzomib (30).

The highest-ranking drug combination with a low magnitude was carfilzomib and LY2157299 (galunisertib) (see Table 4). Galunisertib, a small molecule inhibitor of the transforming growth factor-beta (TGF-β) receptor I kinase, disrupts the phosphorylation of SMAD2, thereby inhibiting the canonical TGF pathway. Given that TGF-β signaling has been implicated in tumor progression and that proteasome inhibition affects cell cycle regulation and apoptosis, this combination theoretically holds great therapeutic benefits (31). However, the potential for overlapping toxicities remains a concern, necessitating further preclinical and clinical research.

Furthermore, the second most-effective combination observed in our ranked list analysis was carfilzomib and selumetinib. Selumetinib, one of the few FDA-approved drugs for NF1, has demonstrated significant efficacy in managing NF1-associated plexiform neurofibromas (PNs) in pediatric patients with inoperable tumors. In a phase 2 clinical trial, selumetinib treatment led to tumor shrinkage and improved clinical outcomes. (6, 32). While the combination of carfilzomib and selumetinib has been lightly studied, their distinct mechanisms—MEK inhibition by selumetinib and proteasome inhibition by carfilzomib (see Figure 9)—suggest that this combination could prove to be increasingly effective. Investigating this combination could provide valuable insights into expanding treatment options for NF1 patients.

**Fig. 9.**
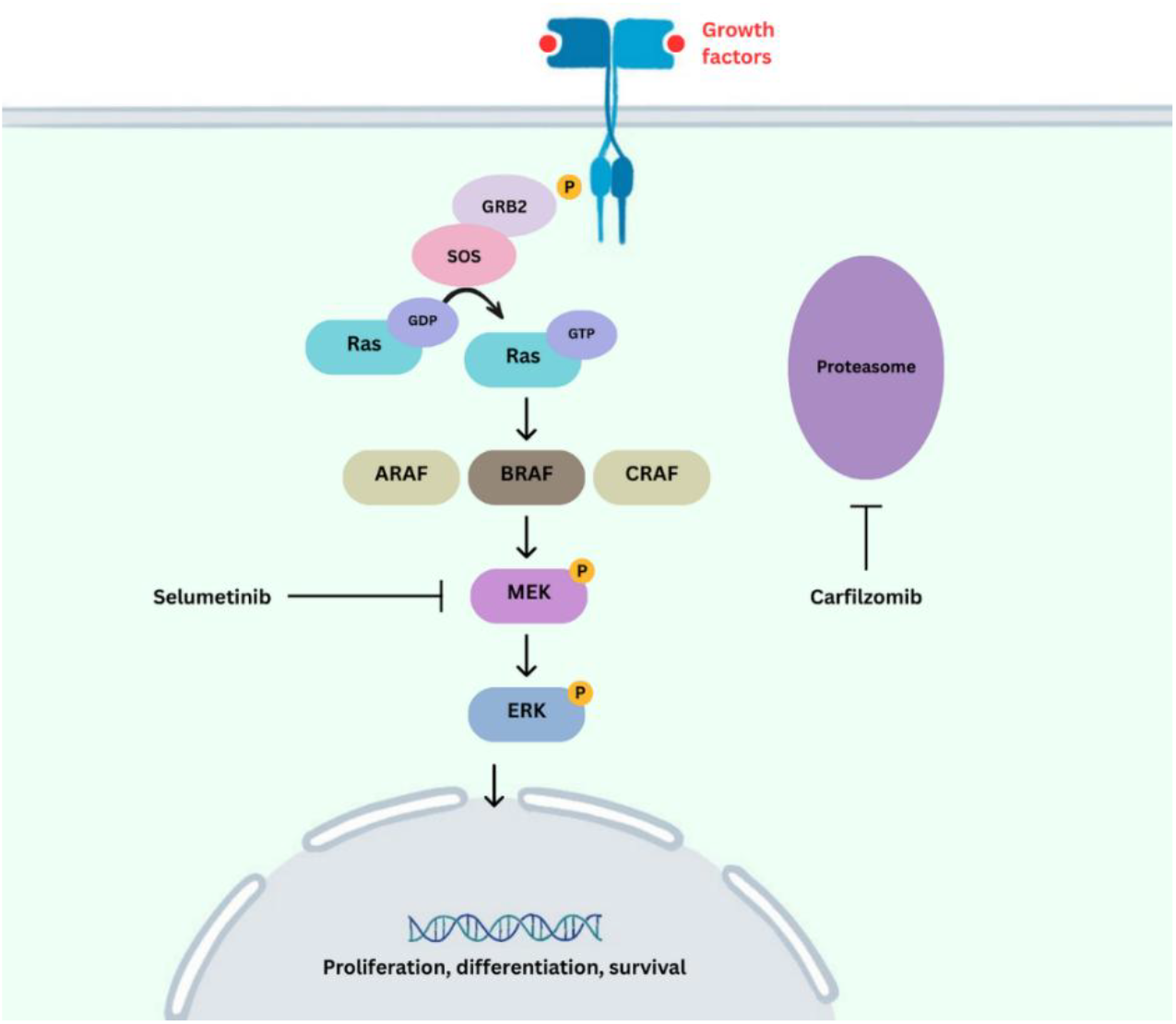
Chart displaying the independent signaling effects of selumetinib + carfilzomib co-therapy. Selumetinib acts as an MEK inhibitor which affects the MAPK signal pathway while carfilzomib acts as a proteasome inhibitor.

Finally, the third highest-ranking combination within Table 4 was carfilzomib and AZD-2014 (vistusertib), an mTOR inhibitor. Though AZD-2014 has been studied in combination with various therapies, information on its interaction with carfilzomib remains scarce. Given the involvement of mTOR signaling in NF1 tumorigenesis, combining an mTOR inhibitor with a proteasome inhibitor, such as carfilzomib, could enhance anti-tumor activity. Further investigation into this combination is necessary to prove its potential efficacy and safety in NF1 treatment.

MEK inhibitors are key components of treatments that aim to repress the mitogen-activated protein kinase (MAPK) signal pathway. This pathway can lead to the proliferation, differentiation, and survival of cancer cells (33). MEK is a family of dual-specific protein kinases that are responsible for the activation of extracellular signal-related kinases (ERKs). MEK itself is activated via serine phosphorylation by kinases such as c-Raf, a-Raf, and b-Raf (33). Selumetinib is a cancer drug that acts as a MEK inhibitor (see Figure 9) and has been especially effective against plexiform neurofibromas which are associated with NF1 (34). In previous trials, it was shown that this treatment reduced tumor size significantly compared to the control group (35). This drug has also shown effectiveness in treating melanoma, differentiated thyroid cancer, and NSCLC (34). Following the success of selumetinib, other MEK inhibitors were formed into new drugs (36). One of these drugs is called mirdametinib and has previously been FDA approved (37). While both drugs have proven to be effective when treating NF1 patients, recent studies have shown that mirdametinib may result in a greater response (38).

### 4.3 Findings in Selumetinib Combinations

Selumetinib and mirdametinib are currently the only two FDA-approved drugs for NF1. Due to the possibility of patients being unresponsive to these drugs, it is crucial to find additional treatments for NF1. The purpose of both this paper’s algorithm and the algorithm in “Integrated Drug Mining Reveals Actionable Strategies Inhibiting Plexiform Neurofibromas” (11) is to facilitate the discovery of potential alternatives to selumetinib and mirdametinib.

Selumetinib ranked relatively high (top 15) in both Sun et. al’s (11); however, in our analysis, there are many drugs that scored better. This resemblance in ranking in the top 15 most effective drugs suggests that selumetinib, while still being an effective treatment for NF1, may not necessarily be the most effective treatment. Additionally, a combination including selumetinib and carfilzomib only appeared once in our top 15 ranked list, emphasizing the potential for more responsive drug combinations. In the ipNF95.6 cell line, the top-ranking selumetinib-based combination was with carfilzomib, a proteasome inhibitor. Other notable pairings included CHS-828 (a NAMPT inhibitor), LY2157299 (a TGF-β receptor I kinase inhibitor), Neratinib (a pan-HER inhibitor), and Tipifarnib (a farnesyltransferase inhibitor).

The mechanisms of action that consistently ranked higher than selumetinib, a MEK inhibitor, includes HSP90 inhibitors, HDAC inhibitors, and mTOR inhibitors. These three inhibitors may be able to target irregularities caused by NF1 in the RAS pathway. A deeper look into these inhibitors may give insight into better treatments than selumetinib.

### 4.4 Comparison to IDAScore Approaches

In contrast to Sun et al. (11), who analyzed drugs individually, our study examined the synergistic effect of drugs. Several drugs from Sun et al.’s study were also top candidates in our analysis. For instance, Sulindac—ranked 6th in Cluster 6 in Sun et al.’s paper—appeared in 26 of the top 100 drug combinations, while tipifarnib, selumetinib, and mirdametinib (PD 0325901) appeared in 33, 35, and 38 of the top 100 combinations respectively. Each drug had a total of 39 combinations.

While Sun et al. focused on identifying drugs compatible with a single compound (selumetinib), our approach added other drug combinations, providing a complementary methodology. Additionally, instead of interpreting combinational effects from single-agent assay data, our study directly evaluated drug combinations using combinational assays, albeit with the limitation of a smaller concentration range. Sun et al. limited their analysis to drugs that are either FDA-approved or in advanced clinical trials (Phase II or III), whereas we included experimental compounds. The top performing drugs identified by our computational methods were almost entirely different from those highlighted by Sun et al., with just a few drugs appearing in both of our studies. Differences in focus and analysis reach could be attributed to these discrepancies, notably in study design; Sun et al. emphasized clinically advanced compounds, while our broader approach included experimental drugs and direct combinational testing, enabling the discovery of novel therapeutic candidates beyond current clinical pipelines.

### 4.5 Disparities, Limitations, and Future Directions

The cell lines used in our dataset from Ferrer et al. (18) utilizes several immortalized cell lines which may exhibit different behavior from in vivo cells. Additionally, the current study uses a limited number of cell lines and could benefit from a larger number of test and reference lines to increase confidence in drug response repeatability. A limited number of cell lines can accentuate inherent drug-response heterogeneity between different cell lines (8, 11) and may have led to the distinct differences in CMRS scores that were noted between the two test cell lines. Moreover, since each drug combination score in this study is based on a single measurement within each matrix, the reliability of the experimental system may have been overestimated (24). Future studies may seek to involve the use of gene regulatory network-based integrated analysis (11) combined with additional reference and test cell lines. While four parameter logistic curve analysis approach are still valid for early drug discovery, they do not align with the exponential costs of large sample size combinational drug studies. In contrast, the CMRS algorithm offers a distinct and more scalable analytical approach. We believe the CMRS algorithm to be a promising foundation to analyze large-scale drug databases targeting combinational drug evaluations.

## Author contributions

Conceptualization: P.O.Z., M.Z., K.Z., S.S., Z.A., O.Z., C.S.

Methodology: P.O.Z., M.Z., S.S., K.Z., Z.A., O.Z., C.S.

Investigation: K.Z., S.S., Z.A., P.O.Z., M.Z., O.Z., C.S.

Visualization: K.Z., Z.A., S.S., P.O.Z., M.Z.

Funding acquisition: M.Z, U.S.

Project administration: K.Z., P.O.Z., M.Z., U.S., F.G.

Supervision: P.O.Z., M.Z.

Writing – original draft: K.Z., S.S., Z.A., O.Z., C.S.

Writing – review & editing: K.Z., S.S., Z.A., O.Z., C.S., P.O.Z., M.Z., F.G.

## Data and materials availability

Quantitative High-Throughput Screening data are available for the MIPE4.0 library of small molecules from https://www.synapse.org/Synapse:syn5611796.

